# Time- and cost-efficient high-throughput transcriptomics enabled by Bulk RNA Barcoding and sequencing

**DOI:** 10.1101/256594

**Authors:** Daniel Alpern, Vincent Gardeux, Julie Russeil, Bart Deplancke

## Abstract

Genome-wide gene expression analyses by RNA sequencing (RNA-seq) have quickly become a standard in molecular biology because of the widespread availability of high throughput sequencing technologies. While powerful, RNA-seq still has several limitations, including the time and cost of library preparation, which makes it difficult to profile many samples simultaneously. To deal with these constraints, the single-cell transcriptomics field has implemented the early multiplexing principle, making the library preparation of hundreds of samples (cells) markedly more affordable. However, the current standard methods for bulk transcriptomics (such as TruSeq Stranded mRNA) remain expensive, and relatively little effort has been invested to develop cheaper, but equally robust methods. Here, we present a novel approach, Bulk RNA Barcoding and sequencing (BRB-seq), that combines the multiplexing-driven cost-effectiveness of a single-cell RNA-seq workflow with the performance of a bulk RNA-seq procedure. BRB-seq produces 3’ enriched cDNA libraries that exhibit similar gene expression quantification to TruSeq and that maintain this quality, also in terms of number of detected differentially expressed genes, even with low quality RNA samples. We show that BRB-seq is about 25 times less expensive than TruSeq, enabling the generation of ready to sequence libraries for up to 192 samples in a day with only 2 hours of hands-on time. We conclude that BRB-seq constitutes a powerful alternative to TruSeq as a standard bulk RNA-seq approach. Moreover, we anticipate that this novel method will eventually replace RT-qPCR-based gene expression screens given its capacity to generate genome-wide transcriptomic data at a cost that is comparable to profiling 4 genes using RT-qPCR.

**‘Software:** We developed a suite of open source tools (BRB-seqTools) to aid with processing BRB-seq data and generating count matrices that are used for further analyses. This suite can perform demultiplexing, generate count/UMI matrices and trim BRB-seq constructs and is freely available at http://github.com/DeplanckeLab/BRB-seqTools

**Highlights:**

- Rapid (~2h hands on time) and low-cost approach to perform transcriptomics on hundreds of RNA samples
- Strand specificity preserved
- Performance: number of detected genes is equal to Illumina TruSeq Stranded mRNA at same sequencing depth
- High capacity: low cost allows increasing the number of biological replicates
- Produces reliable data even with low quality RNA samples (down to RIN value = 2)
- Complete user-friendly sequencing data pre-processing and analysis pipeline allowing result acquisition in a day

## INTRODUCTION

High-throughput sequencing has become the method of choice for genome-wide transcriptomic analyses as its price has substantially decreased over the last years. Nevertheless, the high cost of standard RNA library preparation and the complexity of the underlying data analysis still prevents this approach from becoming as routine as quantitative qPCR, especially when many samples need to be analysed. To rectify overall expenses, the emerging single-cell transcriptomics field implemented the sample barcoding / early multiplexing principle. This reduces both the RNA-seq cost and preparation time by allowing the generation of a single sequencing library that contains multiple distinct samples / cells (Ziegenhain et al., 2017). Such a strategy could also be of value to reduce the cost and processing time of bulk RNA sequencing of large sets of samples (Waszak et al., 2015; Cannavò et al., 2016; Kilpinen et al., 2017; Pradhan et al., 2017). However, there are surprisingly little efforts for explicitly adapting and validating the early stage multiplexing protocols for reliable and cheap profiling of bulk RNA samples.

All RNA-seq library preparation methods are globally relying on the same molecular steps, such as reverse transcription (RT), fragmentation, indexing and amplification. However, when compared side by side, one can observe variation in the order and refinement of these steps (**Figure 1A**). The *de facto* standard workflow (Illumina TruSeq Stranded mRNA) evokes late multiplexing: library indexing at the last amplification step prior to pooling and sequencing, therefore requiring to process all the samples one by one. To overcome this limitation, the RNAtag-seq protocol relies on barcoding of fragmented RNA samples, which allows for early multiplexing and generation of a sequencing library covering the entire transcripts (Shishkin et al., 2015). However, the protocol involving rRNA-depletion and bias-prone RNA adapter ligation (Fuchs et al., 2015) is relatively cumbersome and expensive. Although providing a significantly faster and cheaper alternative, other approaches such as QuantSeq (Lexogen) and LM-seq still require handling every sample individually (Hou et al., 2015) (**Figure 1A)**. In contrast, the early multiplexing protocols designed for single-cell RNA profiling (CEL-seq2, SCRB-seq and STRT-seq) provide a great capacity for transforming large sets of samples, into a unique sequencing library (Hashimshony et al., 2016; Islam et al., 2012; Soumillon et al., 2014). This is achieved by introducing a sample-specific barcode during the RT reaction using a 6-8 nt tag carried by either the oligo-dT or the template switch oligo (TSO). After individual samples have been labeled, they are pooled together and the remaining steps are performed in bulk, thus shortening the time and cost of library preparation. Since the label is introduced to the terminal part of the transcript prior to fragmentation, the reads solely cover the 3’ or 5’ end of the transcripts. Therefore, the principal limitation of this group of methods is the incapacity to address splicing, fusion genes, or RNA editing-related research questions, or to detect allelic imbalance. However, they are proven robust for assessing gene expression levels and identifying differentially expressed (DE) genes. Moreover, most transcriptomics studies do not require or exploit full transcript information, which indicates that standard RNA-seq methods generate more information than typically required, therefore unnecessarily inflating the overall experimental cost.

**Figure 1.**
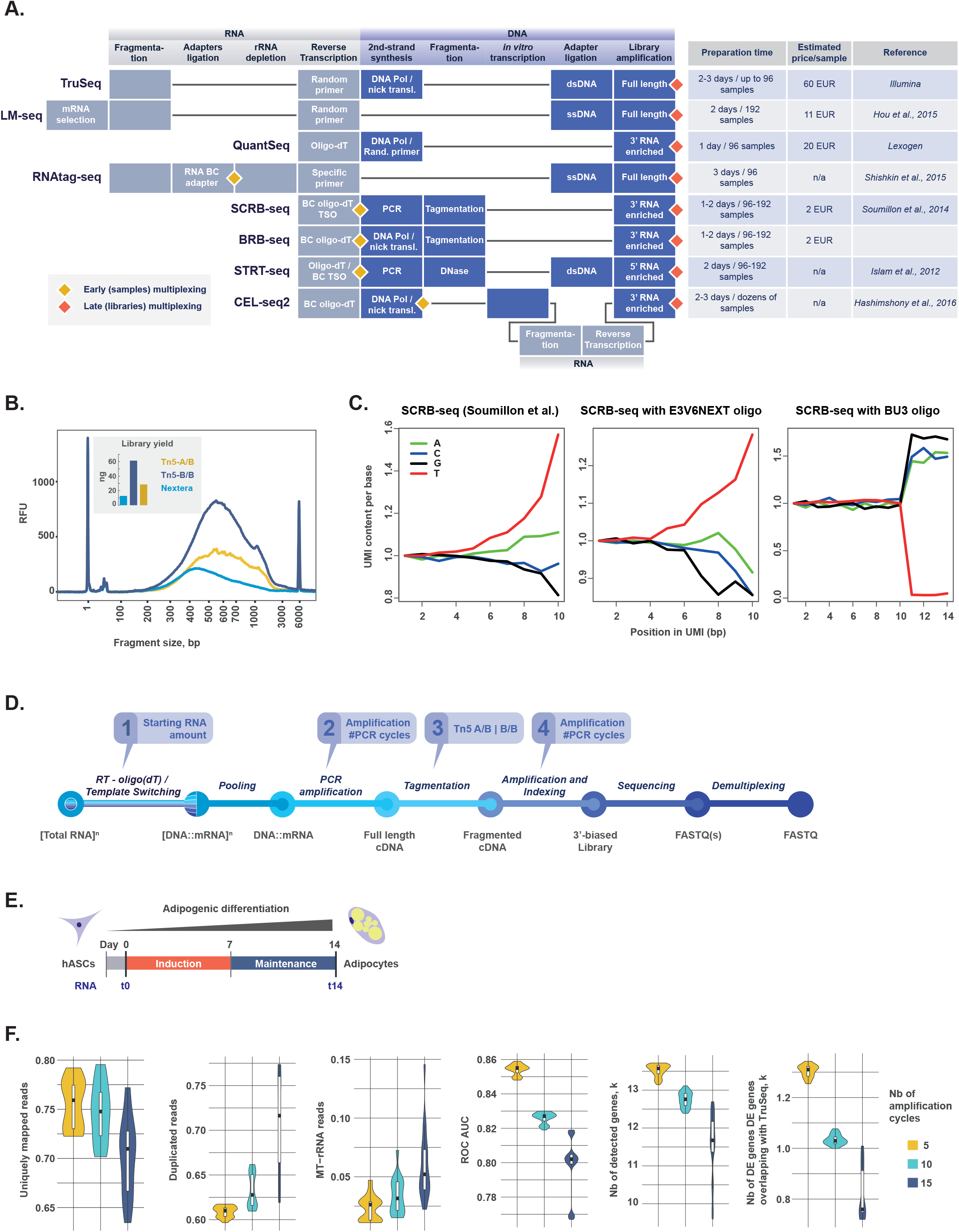
**A.** State-of-the-art bulk and single-cell RNA-seq approaches allowing different levels of samples multiplexing. Every box corresponds to a standardized step of the protocol. Diamonds represent the multiplexing steps: early (yellow) or late (red). The right table shows the throughput of each protocol and the estimated library preparation cost per sample in Euros. BC, barcoded; TSO, template switch oligo. **B and C.** First optimization of the SCRB-seq protocol which led to the novel BRB-seq protocol. **B.** The library profiles and yields after tagmentation with various Tn5 enzymes. **C**. The UMI base content in i) SCRB-seq samples (here D1T0A sample) with E3V6NEXT 10bp UMI oligo from (Soumillon et al., 2014), ii) SCRB-seq performed with bulk RNA and E3V6NEXT oligo, and iii) SCRB-seq with our modified BU3 oligo, bearing a 15bp UMI. **D.** The key steps of the BRB-seq protocol, with four of these considered critical for optimization: i) the amount of input RNA, ii) the number of pre-amplification cycles post RT, iii) the Tn5 enzyme type, and iv) the number of amplification cycles post tagmentation. **E.** The experimental design for preparation of the RNA samples. Human adipocyte stromal cells (hASCs) were differentiated using the adipogenic induction and maintenance cocktails during 7 days each to obtain adipocytes. The cells at both differentiation time points (t0 and t14) were used for RNA extraction. To prepare the libraries, two aliquots of each of two RNA samples at different dilutions (1 – 2000 ng) were used as technical replicates. **F.** The impact of PCR amplification cycles used for the synthesis of full-length dscDNA on the quality of sequencing data, assessed by the percentage of uniquely mapped, duplicated and MT-rRNA reads, Area Under ROC Curve, total number of detected genes and number of DE genes overlapping those detected using TruSeq. For an unbiased comparison, all libraries were randomly downsampled to one million single-end reads (see **Methods**).

In this study, we set out to generate a protocol for bulk RNA profiling that combines the high throughput capacity of single-cell transcriptomics with the high performance of bulk RNA-seq protocols. Inspired by the SCRB-seq method (Soumillon et al., 2014), we developed a novel Bulk RNA Barcoding sequencing (BRB-seq) approach. We have tested and optimized the key steps of the protocol including initial RNA amount, barcoded primer design, number of amplification cycles and tagmentation strategies. We further assessed the performance of BRB-seq relative to Illumina TruSeq Stranded mRNA, the standard for analysing bulk RNA samples, and found that BRB-seq is highly reliable for all assessed quality markers and displays high performance even on fragmented RNA samples.

## RESULTS

### Adaptation of single-cell RNA-seq library preparation method for bulk samples

We selected the SCRB-seq workflow as our starting point to develop the bulk RNA-seq protocol because we judged it to be the most time- and cost-effective amongst the other methods (**Figure 1A**) (Soumillon et al., 2014). SCRB-seq labels each RNA sample with a barcode, introduced with the oligo-dT during template switch assisted RT (**Figure 1A** and **2A**). Moreover, next to the 6 nt barcode, the oligo-dT carries a unique molecular identifier (UMI), a sequence of 10 random nucleotides that allows counting the individual transcripts and evaluating the bias introduced by PCR amplification (Islam et al., 2014; Kivioja et al., 2011). After first strand synthesis, all samples are pooled together and the second strand is generated by PCR to produce full-length double stranded transcripts. The sequencing library is prepared within a two-step process: the cDNA is tagmented (fragmented and tagged) by Tn5 transposase (Picelli et al., 2014) after which the library is enriched by limited-cycle PCR with Illumina compatible adapters. Importantly, this workflow allows to preserve strand specific information.

**Figure 2.**
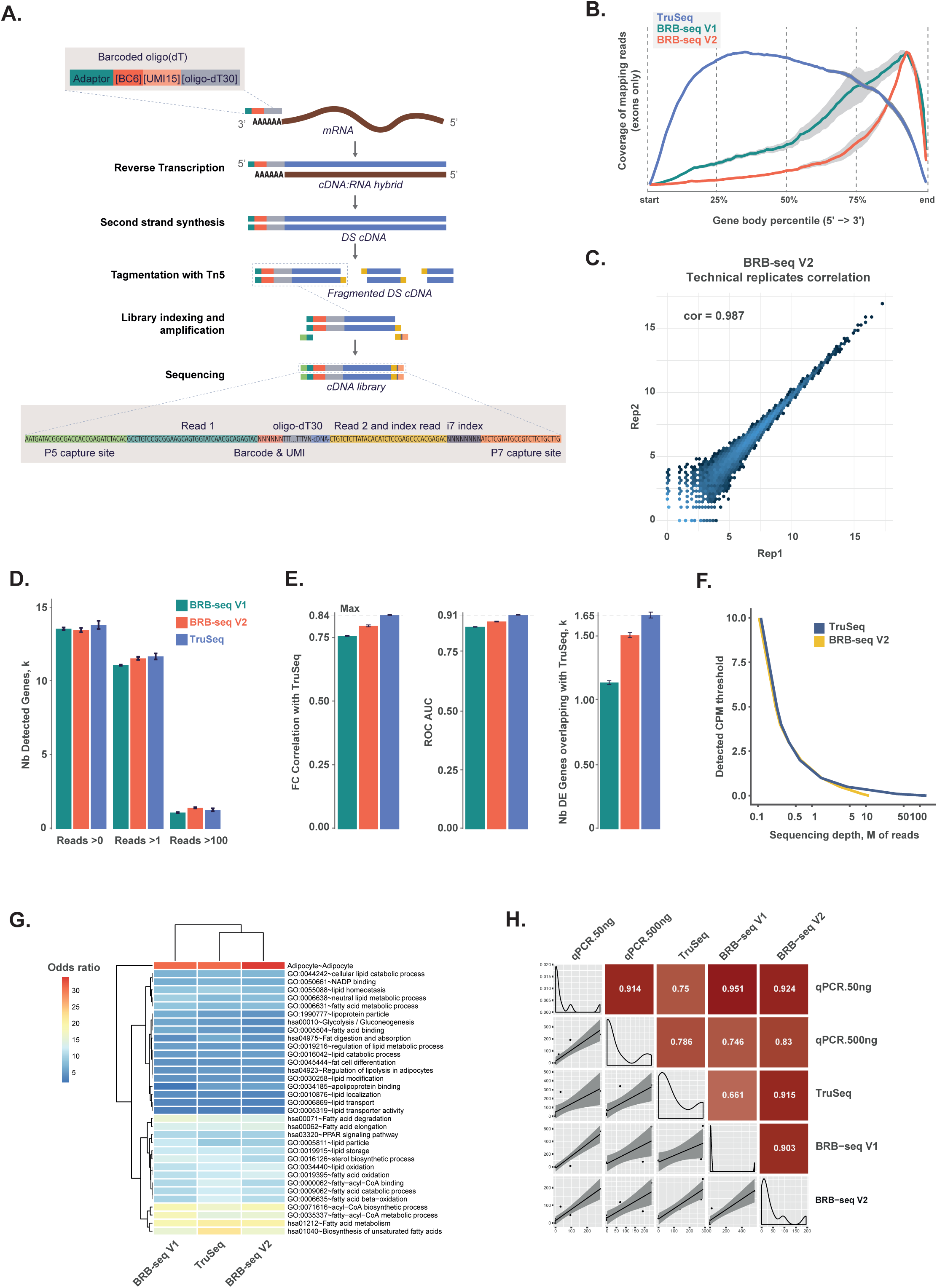
**A.** The final version of the BRB-seq V2 workflow, with second-strand synthesis by nick translation that replaces the PCR amplification in BRB-seq V1. **B.** Read coverage across the gene body for TruSeq, and both versions of BRB-seq. **C.** Correlation of log2 read counts between technical replicates (t14a and t14b) for the BRB-seq V2 workflow (Pearson correlation = 0.987, other methods are compared in **Figure S2D**). **D. & E.** For an unbiased comparison, all libraries were randomly downsampled to one million single-end reads (see **Methods**). **D.** The number of detected genes for TruSeq, and both versions of BRB-seq at different cutoffs. For example, ‘Reads >0’ means that a gene is considered detected if it is covered by at least one read for the given sample. **E.** The comparison of TruSeq, and both versions of the BRB-seq workflow, as represented by three different quality markers (fold change correlation with TruSeq, ROC AUC, and number of DE genes overlapping with TruSeq). **F.** The sequencing depth required for detecting genes with a given CPM expression level using TruSeq and BRB-seq libraries. A sequencing depth is considered sufficient if the gene is detected more than 95% of the time. **G.** The functional enrichment of the identified differentially expressed genes (FC > 2 & FDR < 5%). The odds ratio is calculated using a Fisher’s Exact Test. **H.** Correlation of expression values determined by qPCR (in replicates, with 50ng and 500ng of total RNA used per RT), TruSeq and BRB-seq. Nine target genes with variable expression levels were used for the estimation (see Methods).

As a first step, we designed an experiment using the original primers and reagents of SCRB-seq to test and further optimize its performance with bulk RNA samples. Since we aimed to reduce the cost of library preparation, we also compared the tagmentation efficiency of the Nextera DNA kit versus Tn5 enzymes that were produced in-house following the protocol of (Picelli et al., 2014). The Nextera transposase complex contains a mix of Tn5 enzymes with two different adapter sequences (Tn5-A/B) intended to append either i5 or i7 Illumina indexes to generate compatible sequencing libraries. However, since the SCRB-seq libraries are amplified using only the i7 adapter (and a custom P5-TSO, bearing a P5 capture sequence), the cDNA fragments that are tagmented with the Tn5 that introduces the i5 compatible adapter sequence are not amplified by the limited cycle PCR due to suppression PCR and are thus lost (Siebert et al., 1995). To reduce the loss of cDNA fragments during the tagmentation step, we used Tn5 enzymes loaded with only i7 compatible adapters (Tn5-B/B). This resulted in increased library yield as compared to homemade Tn5 bearing both adapters (Tn5-A/B) or the Nextera (**Figure 1B**).

We further noticed that some UMIs were significantly overrepresented in the library sequencing data. Indeed, UMI read base distribution showed a clear enrichment bias towards UMIs carrying stretches of deoxythymidines (dT) (**Figure 1C**). More specifically, we observed increased occurrences of “T” bases in the UMI sequence in the proximity of the dT stretch (**Figure 1C, left and center panels**). We reasoned that since the stretch of 30 dT is not separated from the UMI sequence in the E3V6NEXT oligo-dT primer, oligonucleotides with longer dT have a higher affinity to the poly-A RNA tail, thus potentially affecting the diversity of the reads. We designed BU3 primers with UMI and oligo-dT sequences separated by five random non-T nucleotides (“V”), therefore increasing the total UMI length to 15 nt (10 “N” + 5 “V”). This proved to be sufficient to reduce the overrepresentation of “T”-containing UMIs (**Figure 1C**, right panel).

### Bulk RNA barcoding and sequencing

To optimize the protocol for bulk RNA samples, we set out to independently evaluate each key step that we deemed critical for the quality of the libraries and resulting sequencing data: i) the amount of input RNA, ii) the number of pre-amplification cycles post RT, iii) the Tn5 enzyme type, and iv) the number of amplification cycles post tagmentation (**Figure 1D**, boxes 1-4). As shown below, these optimization efforts resulted in several important modifications. Therefore, we termed our novel workflow BRB-seq, for bulk RNA barcoding and sequencing, to distinguish it unambiguously from the single-cell-focused SCRB-seq.

For a systematic evaluation of the workflow, we established a set of quality/performance markers to address the alignment efficiency (uniquely mapped reads, mitochondrial-mapping reads), the read diversity (number of detected genes), and more importantly the biological inference that is the main goal of most RNA-seq studies: number and relevance of uncovered DE genes. We employed a reference “gold standard” dataset, generated with the Illumina TruSeq Stranded mRNA protocol using the same RNA samples to estimate the relevance of discovered DE genes. For this, we used samples of total RNA isolated from human adipose stromal cells (hASCs) at two timepoints of adipogenic differentiation: t0 and t14 (non-differentiated ASCs and adipocytes, respectively) with two technical replicates each (**Figure 1E**). To perform an unbiased comparison between the two approaches, the corresponding sequencing libraries were downsampled (DS) to one million single-end reads (see **Methods**).

The data from BRB-seq libraries showed considerable performance variation depending on the used conditions, which could mainly be attributed to the extent of the pre-amplification of full length cDNA (**Figure 1F, S1A-B**). Indeed, libraries with higher amplification cycles yielded an overall lower percentage of aligned reads, higher duplicates rate, less detected genes, and more reads mapping to mitochondrial ribosome genes (MT-rRNA). Similarly, in terms of biological interpretation, the more a sample was amplified, the less DE genes were overlapping with the TruSeq-derived ones. This effect was robustly maintained after examining more detailed performance metrics such as the Area Under the ROC Curve (ROC AUC) that represent detection capabilities of the method across multiple statistical thresholds (**Figure 1F**).

Contrary to our expectations, increasing the amount of input RNA showed only a negligible performance improvement in terms of number of overlapping DE genes. This indicates that the BRB-seq protocol works similarly good over a large range of input RNA, 10 - 2000 ng of RNA per sample (**Figure S1A**). Since the cDNA prepared from as little as 4 ng of total RNA should have been amplified for at least 15 PCR cycles to obtain a sufficiently high cDNA amount that would be compatible with library preparation, we were not able to test different pre-amplification conditions for the library containing 1 ng of input RNA per sample. However, the overall performance of this library was comparable to those with higher input RNA amounts that were amplified for 15 cycles (**Figure S1A**). Thus, using as low as 1 ng of RNA for library preparation produces data that is of similar quality to that of libraries derived from greater RNA amounts.

The impact of the other examined factors that could introduce variation when generating full-length cDNA was found to be relatively minor. For example, increasing the amount of cDNA used for tagmentation did not dramatically affect the detection of DE genes (data not shown). Interestingly, the use of Tn5 with different preloading adapter pairs (**Figure S1B**), and post-tagmentation library amplification (**Figure S1C**) had a visible but marginal effect on overall performance.

### Second-strand synthesis without amplification improves data quality and biological relevance

Our results revealed an inverse correlation between the complexity/diversity of BRB-seq libraries and the number of PCR amplification cycles that were used to generate full-length double stranded cDNA. We concluded that the method could be improved by using the modified Okayama and Berg procedure (Gubler and Hoffman, 1983) to generate double stranded cDNA instead of PCR amplification. Importantly, since this method relies on second strand generation via RNA primer-depended nick translation by DNA polymerase I, there is no need to include the template switch oligo in the first strand synthesis (**Figure 2A**, step 2).

For the second strand synthesis by nick translation (SSS) reaction, we used NEB enzymes that were reported to work optimally with first-strand cDNA produced by SuperScript II (SSII) RT. This modification of the protocol led us to its two main versions. The initial version (BRB-seq V1), synthetizes the first strand using Maxima Minus H (MMH) and the second strand by PCR. The new version (V2), involves SSII and SSS respectively (results on all possible enzyme/second strand synthesis combinations can be found in **Figure S2A-C**).

The yield of full-length cDNA generated with BRB-seq V2 was lower compared to that obtained with V1 (**Figure S2A**). However, the new workflow proved to be reliable to produce libraries in sufficient amounts and overall quality for sequencing. As a first QC, we compared the read coverage over exonic regions, and found that V2 libraries were more concentrated at the 3’ end of transcripts than those produced by V1 (**Figure 2B**). This broader read coverage of the V1 protocol can be explained by the higher bias and rate of MT-rRNA reads in libraries prepared with the MMH enzyme (**Figure S2B**), pointing to a previously reported higher enzymatic activity of the latter compared to SSII (Bagnoli et al., 2017).

Another improvement of BRB-seq V2 was an increased ratio of reads mapping to annotated genes, namely ~76%, compared to ~65% produced with BRB-seq V1 (**Figure S2C**). This result is much closer to the overall proportion obtained by TruSeq (~83-84%) and highlights the fact that BRB-seq V2 libraries exhibit less bias/noise resulting from adapter and polyA contamination. Moreover, like TruSeq, the V2 libraries produced highly correlated technical replicates, highlighting its robustness and analytical sensitivity (**Figure 2C** and **Figure S2D**). Similarly, the V2 libraries reads displayed greater diversity allowing to detect a similar number of genes than TruSeq at the same sequencing depth (**Figure 2D** and **Figure S2E**). Further analyses revealed the higher accuracy of the improved V2 workflow at recovering relevant biological signal compared to TruSeq, represented by the increased fold change correlation, AUC values (ROC curve), and number of DE genes overlapping with TruSeq (**Figure 2E** and **Figure S2F**).

Comparison of the initial TruSeq library that we reference as “gold standard” (~30 millions paired-end reads) with its downsampled counterpart (one million single-end reads) displayed an 84% correlation, which sets a threshold of maximum attainable accuracy at this sequencing depth (**Figure 2E** and **Figure S2F**). After this ~30-fold downsampling, we identified 1656 DE genes, representing 36% of all “gold standard” DE genes (4566 total). Moreover, the DE genes that were not detected tend to be lowly expressed (**Figure S2G**). We therefore conclude that high sequencing depth (> 5 M reads) is not essential to reliably identify DE genes with medium or high expression levels (CPM > 1). However, higher coverage is needed to profile lowly expressed genes (**Figure 2F**). Importantly, we do not observe a strong difference in detection capacity between BRB-seq and TruSeq across the same sequencing depths (**Figure 2F**).

We further investigated whether DE genes discovered with the different methods were biologically relevant. For this, we conducted a functional enrichment of the DE genes that were up-regulated in the differentiated hASC cells using adipocyte-related gene sets from KEGG, Gene Ontology (GO), and Gene Atlas databases. Overall, both BRB-seq V1 and V2 DE genes were strongly enriched in these adipocyte gene sets (**Figure 2G**). Nevertheless, the data obtained with the V2 protocol clustered closer to TruSeq than BRB-seq V1. It is also worth noting that the “Adipocyte” gene set (from Gene Atlas database) was slightly more enriched with BRB-seq V2 as compared to TruSeq at similar sequencing depth (**Figure 2G**). Taken together, this confirms the greater overall performance of the V2 protocol and we therefore define this version as the final BRB-seq workflow (**Figure 2A**).

### Functional validation of BRB-seq

Given the performance of BRB-seq, combined with the fact that it is fast and cost-efficient, we envisioned that it could potentially become an alternative to RT-qPCR assays, especially when large sets of samples need to be profiled. To confirm that BRB-seq libraries can produce reliable gene expression results, we compared it to RT-qPCR data. We evaluated 9 genes that are expressed at different levels in adipocytes. We performed two RT-qPCR replicates, one with 50 ng of RNA and the other with 500 ng. The same RNA sample was used to prepare first strand reactions for BRB-seq and TruSeq libraries. After normalization to *HPRT1* expression, we assessed the correlation of expression values between each of the methods (**Figure 2H**). We observed that both BRB-seq and TruSeq highly correlate with qPCR (~0.8-0.9 correlation values). Interestingly, BRB-seq correlates even slightly better with qPCR than TruSeq, and this effect was observed for both qPCR replicates.

### BRB-seq application for degraded RNA samples

It is well established that the TruSeq Stranded mRNA method performs poorly on degraded RNA samples given the intrinsic requirement of this method to have an RNA Quality Number (equal to RIN, RNA Integrity Number) ≥ 7-8. This may reflect the fact that full-length transcripts are sequenced, thus requiring high-quality, intact RNA for accurate detection and quantification. Since 3’ RNA fragment quantification is known to be a robust way to estimate differential gene expression in samples with low RQNs (Sigurgeirsson et al., 2014), we decided to evaluate the performance of BRB-seq on fragmented RNA samples with low RQN values. For this, we employed chemical RNA fragmentation by incubation at 65°C in the presence of Mg^++^ cations for one or two minutes, which resulted in a significant reduction in overall RNA size and in a drop of RQN values (**Figure S3A**). Specifically, we observed an RQN of 9.8 and 8.9 for non-degraded samples, 4.7 and 6.4 for samples subjected to one minute of fragmentation and down to 2.2 and 4.6 after two minutes of processing for t0 and t14 respectively. We used these three time points to prepare BRB-seq libraries (V2) using 100 ng of each RNA sample.

We observed a clear trend indicating a reduction in overall gene expression correlation (with data from non-degraded samples) with declining RQN values for each individual RNA sample. However, this correlation remained above 97%, even for samples with very low RQN (**Figure 3A**), indicating a high performance of the method, independent of RNA quality. We investigated further the impact that fragmentation has on the biological relevance of the genes that were detected with respect to characterizing hASC differentiation. As expected, we observed a clear inverse correlation between the quality of the results and the sample RQN values. Yet, the fold change correlation and ROC AUC values between the fragmented and intact samples were maintained at high levels (**Figure 3B**). The number of detected DE genes decreased in highly fragmented samples, however most of the DE genes could still be recovered. Together, these data show that BRB-seq allows reliable differential gene expression and functional enrichment analyses using RNA samples with very low RQN values.

**Figure 3.**
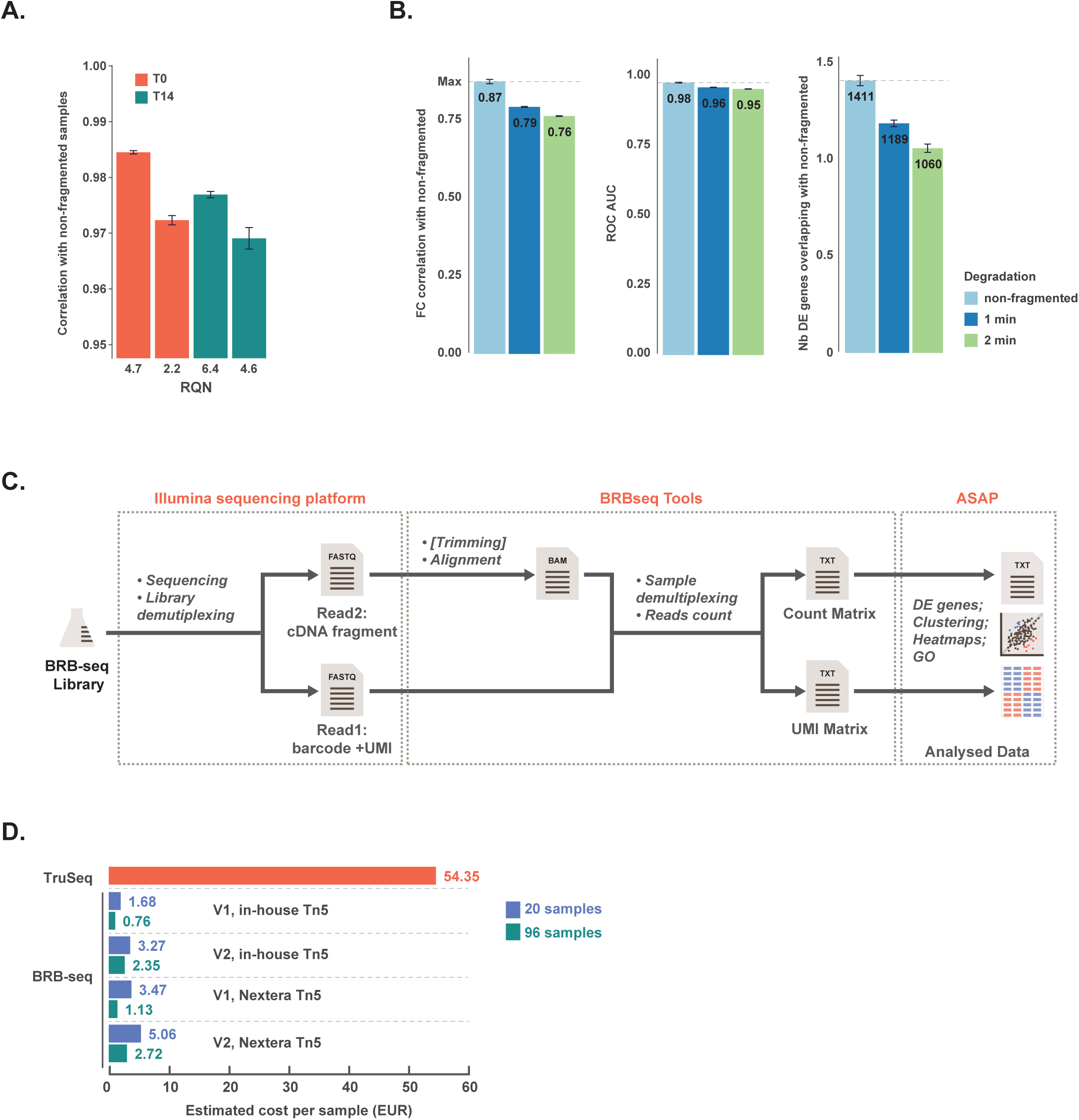
**A. & B.** For an unbiased comparison, all libraries were randomly downsampled to one million single-end reads (see **Methods**). **A.** Correlation (Pearson) between log2 read counts of intact versus fragmented (RNA quality number (RQN)=8.9 & 9.8 for T0 and T14 respectively) samples. **B.** Comparison of intact RNA samples versus their degraded counterparts after one or two minutes of fragmentation, showing three different quality markers (fold change correlation, ROC AUC, and number of DE genes overlapping with non-fragmented samples). **C.** Schematic representation of the BRB-seq library post-sequencing data processing pipeline. It includes the BRB-seqTools module (available on github) that can perform reads trimming (optional) and alignment, sample demultiplexing and generation of a count table. The count table can be further analyzed by standard algorithms or loaded into ASAP, a web-based analytical interface that facilitates data exploration and visualization (Gardeux et al., 2017). **D.** The estimated cost of library preparation per sample for TruSeq and both versions of BRB-seq. The final cost depends on the number of samples that are pooled together and whether in-house or commercial Nextera Tn5 is used.

### BRB-seq data analysis pipeline and considerations

Upon sequencing of the BRB-seq libraries, highly multiplexed datasets are produced which may pose analytical problems, specifically for users with limited bioinformatics skills. To make the entire workflow of the method accessible to end users who lack a strong computational background, we aimed at simplifying the analysis of BRB-seq sequencing data. For this, we developed a complete tool suite (http://github.com/DeplanckeLab/BRB-seqTools) allowing to perform all the required post-sequencing tasks up until the generation of the read/UMI count matrix (**Figure 3C**). Thereafter, the data can be processed with conventional R scripts/packages to perform the required analyses. Alternatively, the count matrix file can be supplied to ASAP (https://asap.epfl.ch/), a web-based platform devoted to comprehensive transcriptome analyses and developed in our lab (Gardeux et al., 2017). In addition to demultiplexing and generating the count/UMI matrix, the BRB-seqTools suite also allows the user to perform specific read trimming to remove sequences that are produced because of BU3 primer and polyT contamination (5 to 10% of raw sequencing reads). Even though such trimming does not significantly change the overall downstream data quality (**Figure S3B**), it does highly enhance overall read quality, which we judge to be of analytical importance. Consequently, along with the method itself, we provide a seamless pre- and post-treatment pipeline for enabling any user to perform a state-of-the-art analysis of their BRB-seq data.

## DISCUSSION

We developed BRB-seq, a bulk RNA barcoding and sequencing approach, that enables genome-wide gene expression analyses of hundreds of RNA samples simultaneously, in an effort- and cost-efficient manner. In our experience, following the BRB-seq workflow, one person can easily prepare sequencing libraries containing 192 samples within 7 - 8 hours given that the projected hands-on time is around 2 hours. The number of samples in one library is scalable and merely depends on the number of available barcodes and desired sequencing depth per sample. We used the 96 barcodes from the SCRB-seq protocol, but practically any set of barcodes of different length can be used to generate the BRB-seq adapters. Along with being fast and easily manageable, the protocol’s high advantage is its low cost (down to 2-3 EUR per sample, **Figure 3D**). Adding the sequencing cost, we estimate the total expense to be around 20 EUR per sample, considering that one NextSeq 500 High Output 75 cycles kit run provides enough reads to support a library of 96 human or mouse samples and more if smaller transcriptomes are probed. Thus, this estimation is entirely guided by the type of desired analysis or organism, and by the relative expression of specific genes of interest, which leaves sufficient space for optimization of sequencing depth and hence even greater cost reduction.

The improved BRB-seq workflow with second strand synthesis based on nick translation rather than PCR amplification generated the data with the highest level of similarity to TruSeq, specifically as measured by the number of mappable reads and identified DE genes. Importantly, the first strand synthesis in the final (V2) protocol does not require a template switch oligo, allowing to alleviate the TS-associated bias towards fully reverse transcribed molecules and potential artifacts related to strand invasion (Shapiro et al., 2013; Tang et al., 2013).

The sole inherent limitation of the BRB-seq method is the requirement to accurately assess RNA sample amounts prior to RT as any inter-sample variation will result in uneven distribution of sequencing reads. In our experience, this issue is solved through re-quantification of intermediate RNA dilutions that are prepared to normalize the concentration variations.

Currently, there are only few other RNA-seq approaches that provide comparably cheap or fast solutions for transcriptomics profiling of bulk RNA (**Figure 1A**). These include LM-seq that allows the detection of full-length transcripts but requires oligo-dT purified mRNA for RT using a random primer (Hou et al., 2015). This method is at least five times more expensive than BRB-seq and importantly, the samples are pooled together at the very last step just prior to sequencing. This implies that eventually every sample is processed separately which makes the library preparation workflow almost as demanding as the standard TruSeq Stranded mRNA. The same applies to the commercially available QuantSeq kit (Lexogen) that also relies on late stage multiplexing. Therefore, it remains costly and comparatively cumbersome for processing many samples at once.

The amount of initial RNA used within the range of 10 – 2000 ng per sample does not substantially affect the quality of the data. Therefore, we recommend the total RNA content in the pool to be in the range of 1 – 2 µg to reduce any potential performance variation of the second strand synthesis step. This corresponds roughly to 10 - 20 ng per sample for a library of 96 samples, or 50 - 100 ng when 20 samples need to be mixed in one library. Our data also suggests that an input RNA amount as low as 1 ng might equally produce a reliable library. However, we recommend in this case to pool multiple samples to assure that the dscDNA is of sufficient quantity for tagmentation. As it is sometimes complicated to assess how deep a sample should be sequenced, we also provide an estimation of the number of sequencing reads that are required to detect a particular gene (95% chance of having at least 1 read), given its CPM expression (**Figure 2F**).

We anticipate that BRB-seq may become an attractive option for routine gene expression analysis and ultimately replace large RT-qPCR assays. Assuming that the current cost of one qPCR reaction is in the range of 2 – 3 EUR, the evaluation of three to four target genes in triplicate (~20 qPCR reactions) will cost approximately the same as one full transcriptome analysis produced by BRB-seq (20 EUR). Importantly, low library preparation cost and time implies that more biological replicates can be profiled, which will greatly increase the statistical power underlying any DE analysis.

Similar to the previously shown capacity of 3’ RNA-seq methods to efficiently work with degraded samples (Sigurgeirsson et al., 2014), BRB-seq libraries display a high level of correlation of detected genes between samples with high and very low RQN values. This robustness extends to DE analyses, revealing that BRB-seq is a potentially valuable approach to profile samples with low quality / degraded RNA. Therefore, we anticipate that BRB-seq might prove also useful for the gene expression profiling of challenging clinical samples.

Finally, the BRB-seq protocol still features the UMI concept, which reflects its single-cell origin. However, this may be less useful for bulk RNA-seq data since the higher input RNA amount requires less amplification. Indeed, we observed only a slight increase in sensitivity when using the UMIs (**Figure S3C**). This is in line with similar conclusions which stated that the removal of UMI identical reads only slightly improves FDR (Parekh et al., 2016). Nevertheless, we kept the UMI in the BRB-seq construct to allow the user to overcome the amplification bias when samples with lower RNA quantities (<1 ng) need to be processed. Also, the UMI provides a good way of unbiased estimation of the duplication ratio compared to computationally guided estimates (Picard (http://broadinstitute.github.io/picard/) or fastQC (http://www.bioinformatics.babraham.ac.uk/projects/fastqc)) which tend to inflate duplicate proportions with increasing sequencing depth. It is worth noting that the user can very simply change the oligo and remove the UMI construct, or keep it but not sequence it for lowering costs.

Importantly, we developed a BRB-seq analytical toolbox that together with the protocol provides a holistic solution for gene expression analysis: from the RNA sample to the gene ontology data. This toolbox, empowered by the web-based count table analysis platform (ASAP) (Gardeux et al., 2017), allows to process the sequencing data within a day in a user-friendly manner even when lacking bioinformatics skills to obtain and visualize gene expression data.

## MATERIALS AND METHODS

### Bioethics

The work on human adipocyte stromal cells (hASC) cultures derived from human lipoaspirate samples is approved by the ethical commission of Canton Ticino (CE 2961 from 22.10.2015) and conforms with the guidelines of the 2000 Helsinki declaration. The anonymized samples were collected under signed informed consent.

### Cell culture

hASC were obtained from a fresh lipoaspirate as following. 50 ml of lipoaspirate was washed twice with 40 ml of DPBS Ca+/Mg+ (Gibco, #14040091) in 100 ml syringes and incubated with 0.28 U/ml of liberase TM (Roche, #05401119001(ROC)) for 45 min at 37ºC under agitation. The digested tissue was mixed with 40 ml of CRB (1% human albumin (CSL Behring) in 40 ml of DPBS -/- (Gibco, #14190094)) and shaken vigorously to liberate the stromal cells. The aqueous phase was recovered and centrifuged at 400*g* for 5 minutes at RT. The cell pellet was resuspended in 15 ml of CRB and filtered through a 100 µm and then 40 µm cell strainer to ensure a single cell preparation, centrifuged and resuspended in Minimum Essential Medium (MEM) alpha (Gibco, #32561037) supplemented with 5% human platelet lysate (Cook Regentec, #G34936) and 50 µg/mL Primocin (InvivoGen, #ant-pm-1). hASCs were cultured in the same media composition until 70-80% confluency and detached using TrypLE Select (Life Technology, #1256311) for passaging.

For the adipogenic differentiation, cells at confluence were treated with induction cocktail from Adipogenic BulletKit (Lonza, #PT-3004) for 7 days, followed by treatment with maintenance cocktail for another 7 days.

### RNA samples for libraries preparation

Total RNA was isolated using TRI Reagent (Molecular Research Center, #TR118) and followed by double precipitation with ethanol. The RNA concentration was determined using the Qubit RNA HS Assay Kit (Invitrogen, #Q32852) and integrity was assessed using a Fragment Analyzer (Advanced Analytical). The RNA from each differentiation time point was used in two technical replicates, resulting in four samples pooled per library. Libraries were prepared with the BRB-seq protocol using the amounts of total RNA ranging from 1 ng to 2 µg per sample (**Supplementary Table 1**).

RNA fragmentation was done using the NEBNext Magnesium RNA Fragmentation Module (NEB, #E6150S) with incubation time at 94°C for one or two minutes. This resulted in RNA with a variable extent of degradation and corresponding RQN values.

### RT-qPCR

For RT-qPCR, 50 ng or 500 ng of total RNA was used to generate the first strand using 1 µL of Superscript II (Invitrogen, #18064014) and 1 µL of anchored oligo-dT (ThermoFisher Scientific, #AB1247) in 20 µL total reaction mix following the protocol. cDNA was diluted five times using nuclease free water and 2 μL was used for each qPCR reaction. Quantitative real-time PCR was performed in three technical replicates on the ABI-7900HT Real-Time PCR System (Applied Biosystems) using the PowerUp SYBR Green Master Mix (Applied Biosystems, #A25742) using standard procedures. The qPCR primers for the targets genes (*ADIPOQ, AXIN2, BCAT, CEBPB, FABP4, HPRT, LEP, LPL, PNPLA2 & PPARG*, see **Supplementary Table 2**) were designed with Primer3 software (Untergasser et al., 2012).

### BRB-seq protocol

#### Reverse transcription/template switching

For the initial version of the protocol (V1) various amounts of RNA (50 pg - 2 µg) were reverse transcribed using 0.125 µL of Maxima H Minus Reverse Transcriptase (ThermoFisher Scientific, #EP0753), 2 µL of RT buffer, 1 µL dNTP (0.2mM), 1 µL of 10 µM template switch oligo (TSO, IDT) and 1 µL of 10 µM barcoded oligo-dT (BU3, Microsynth) in a 96 well plate (for the list of oligos used see **Supplementary Table 2**). After addition of RNA, BU3 primers and dNTP, the plate was incubated at 65°C for 5 minutes and then put on ice. The TSO, RT buffer and enzymes were added to each well and the plates were incubated at 45°C for 90 minutes. Alternatively, for the V2 of the protocol the RNA was reverse transcribed using Superscript II (Invitrogen, #180640). After incubation of RNA, BU3 primer and dNTP at 65°C, the following mix was added into each well: 4 µL of 5x First Strand Buffer, 2 µL DTT (0.1M) and 0.25 µL of Superscript II enzyme (Invitrogen, 180640). The reaction was incubated at 42°C during 50 minutes followed by inactivation at 70°C for 15 minutes. After RT, all the wells were pooled together and purified using DNA Clean & Concentrator-5 kit (Zymo Research, #D4014) with 7x DNA binging buffer and single column. After elution with 20 µL of nuclease-free water, the samples were incubated with 1µL Exonuclease I (NEB, #M0293) and 2 µL of 10X reaction buffer at 37°C for 30 minutes, followed by enzyme inactivation at 80°C for 20 minutes.

#### Second-strand synthesis

For the V1 protocol 20 µL of pooled and ExoI treated cDNA was PCR amplified with Advantage 2 Polymerase Mix (Clontech, #639206) in 50 µL total reaction volume using 1 µL of 10 µM LA_oligo (Microsynth) primer 1 µL of dNTP (0.2mM), 1 µL of Advantage 2 polymerase mix, 5 µL of Advantage 2 PCR buffer and 22 µL of water following the program (95°C – 1 minute, 10 cycles: 95°C – 15 seconds 65°C – 30 seconds, 68°C – 6 minutes; final elongation at 72°C – 10 minutes). Full-length double stranded cDNAs were purified with 30 µL (0.6X) of AMPure XP magnetic beads (Beckman Coulter, #A63881) and eluted in 20 µL of water. In the V2 protocol the second stand was synthetized following the modified Okayama and Berg method (Gubler and Hoffman, 1983). For that the mix containing 2 µL of RNAse H (NEB, #M0297S), 1 µL of E. coli DNA ligase (NEB, #M0205L), 5 µL of E. coli DNA Polymerase (NEB, #M0209L), 1 µL of dNTP (0.2mM), 10 µL of 5X Second Stand Buffer (100mM Tris-HCl (pH6.9) (AppliChem, #A3452); 25 mM MgCl2 (Sigma, #M2670); 450 mM KCl (AppliChem, #A2939); 0.8 mM β-NAD; 60 mM (NH4)2SO4 (Fisher Scientific Acros, #AC20587) and 11 µL of water was added to 20 µL of cDNA after exonuclease treatment on ice. The reaction was incubated at 16°C for at least 2,5 hours or overnight. The product was purified with AMPure XP magnetic beads (Beckman Coulter, #A63881) and eluted in 20 µL of water.

#### Library preparation and sequencing

The sequencing libraries were prepared by tagmentation of 1-50 ng of full-length double stranded cDNA. Tagmentation was done with either Illumina Nextera XT kit (Illumina, #FC-131-1024) following the manufacturer’s recommendations or with in-house produced Tn5 preloaded with dual (Tn5-A/B) or same adapters (Tn5-B/B) under the following conditions: 1 µL (11 µM) Tn5, 4 µL of 5x TAPS buffer (50 mM TAPS (Sigma, #T5130), 25 mM MgCl_2_ (Sigma, #M2670)) in 20 µL total volume. The reaction was incubated 10 minutes at 55°C followed by purification with DNA Clean & Concentrator-5 kit (Zymo Research) and elution in 21 µL of water. After that tagmended library (20 µL) was PCR amplified using 25 µL NEBNext High-Fidelity 2X PCR Master Mix (NEB, #M0541L), 2.5 µL of P5_BRB primer (5 µM, Microsynth) and 2.5 µL of oligo bearing Illumina index (Idx7N5 5µM, IDT) using the following program: incubation 72°C — 3 mins, denaturation 98°C — 30 seconds; 10 cycles: 98°C — 10 seconds, 63°C — 30 seconds, 72°C — 30 seconds; final elongation at 72°C – 5 minutes. The fragments ranging 200 - 1000 bp were size-selected using AMPure beads (Beckman Coulter, #A63881) (1st round 0.5X beads, 2nd - 0.7X). The libraries were profiled with High Sensitivity NGS Fragment Analysis Kit (Advanced Analytical, #DNF-474) and measured with Qubit dsDNA HS Assay Kit (Invitrogen, #Q32851) prior to pooling and sequencing using the Illumina NextSeq 500 platform using custom ReadOne primer (IDT) and the High Output v2 kit (75 cycles) (Illumina, #FC-404-2005). The library loading concentration was 2.2 nM. The read1 sequencing was performed for 6 – 21 cycles and read2 for 54 −70 cycles depending on the experiment.

### RNA library preparation with TruSeq

TruSeq libraries were prepared with 1 µg of total RNA using the TruSeq Stranded mRNA Library Prep Kit (Illumina, #RS-122-2101) and following the manufacturer's instructions. Four libraries were paired-end sequenced (75 nt each) with the NextSeq 500 using the Mid Output v2 kit (150 cycles) (Illumina, #FC-404-2001).

### Pre-processing of the data – demultiplexing and alignment

The raw reads from BRB-seq experiments carry two barcodes, corresponding to the late and early step multiplexing. The late step multiplexing using Illumina indexes is common to standard protocols and is used to separate the libraries. The early barcode is specific to the BRB-seq protocol and is used to separate the multiplexed samples from the bulk data. The first demultiplexing step was performed by the sequencing facility using bcl2fastq software. Then, the data consists of two FASTQ files (R1 and R2). The R2 FASTQ file was aligned to the Ensembl r87 gene annotation of the hg38 genome using STAR (Version 2.5.3a) (Dobin et al., 2013), with default parameters prior to the second demultiplexing step. Then, using the BRB-seqTools suite (available at http://github.com/DeplanckeLab/BRB-seqTools), we performed simultaneously the second demultiplexing, and the count of reads/transcripts (UMI) per gene from the R1 FASTQ and the aligned R2 BAM files. This generated two count matrices (reads and UMI) that were used for further analyses. In parallel, we also used the BRB-seqTools suite for demultiplexing the R1/R2 FASTQ files and producing one FASTQ file per sample. This was required for being able to generate the downsampling of every sample. In this case, FASTQ files were aligned using STAR and HTSeq (Version 0.9.1) (Anders et al., 2015) was used for producing the count matrices.

### mRNA-seq computational analysis and detection of DE genes

All downstream analyses were performed using R (Version 3.3.1, https://cran.r-project.org/). Library normalization and expression differences between samples were quantified using the DESeq2 package (Love et al., 2014), with cutoff of |FC| ≥ 2 and FDR ≤ 0.05. Further functional enrichments were performed using Fisher’s Exact Test on Gene Ontology (Ashburner et al., 2000), KEGG (Kanehisa and Goto, 2000), and Gene Atlas (http://www.genatlas.org/) databases.

### Downsampling of TruSeq and BRB-seq samples

All samples were randomly downsampled to different levels as indicated for every case. To avoid transferring alignment-related issues to the downstream analyses, we did not downsample at the level of the FASTQ files. Indeed, to be able to keep some information about the reads before their mapping to genes (such as duplicates or UMI), we chose to perform the downsampling at the level of the BAM files, just before performing the htseq-count step. For reproducibility and robustness of the results, we chose to generate 5 downsampled BAM for each replicate (i.e. 20 downsamples per library, since every library contains 4 samples originally).

### TruSeq and BRBseq comparison

Coverage over the gene body was computed using the RSeQC suite v.2.6.1 (Wang et al., 2012) with the geneBody_coverage.py script. We used the full list of genes from the hg38 assembly provided on the software web page. ROC and Precision Recall (PR) curves were produced using the set of 4566 DE genes identified using full paired-end TruSeq samples with the DESeq2 package. This set represents a self-assigned “gold standard”, i.e. the positive set, while the negative set constitutes of all genes expressed as detected by TruSeq but not identified as DE. Then, for every comparison, we applied DESeq2 and used the full list of ranked p-values to compare to the “gold standard”. False Positive Rate, True Positive Rate and precision (for Precision-Recall and ROC curves) were computed for every p-value cutoff of the ranked p-value list, thus generating the curves. Area under the curves were computed using the *rollmean* function of the *zoo* package in R. Mitochondrial RNA content (called MT-rRNA content in the figures) was assessed using only two MT-rRNA genes that are known to be the main representatives of any mitochondrial contamination: *MT-RNR1* and *MT-RNR2*.

### Data access

All the datasets have been submitted to ArrayExpress under accession number E-MTAB-6469.

## ACKNOWLEDGEMENTS

We thank all the members of Deplancke’s lab and particularly Johannes Bues for useful discussions. We thank Antonio Meireles Filho and Romane Breysse for sharing the in-house produced Tn5. We thank Jo Vandesompele, Petra C. Schwalie, Wanze Chen and Bastien Mangeat for their comments on the manuscript. We also thank Bastien Mangeat for his assistance in library preparation and sequencing. We thank the VITAL-IT infrastructure that supported our computational analyses.

## Author Contributions

DA, VG and BD conceived and planned the study and prepared the manuscript. DA planned and performed the experimental work and along with JR performed the library preparations. VG performed the RNA-seq computational analyses. All authors discussed the results and commented on the paper.

---

**Supp Figure S1.**
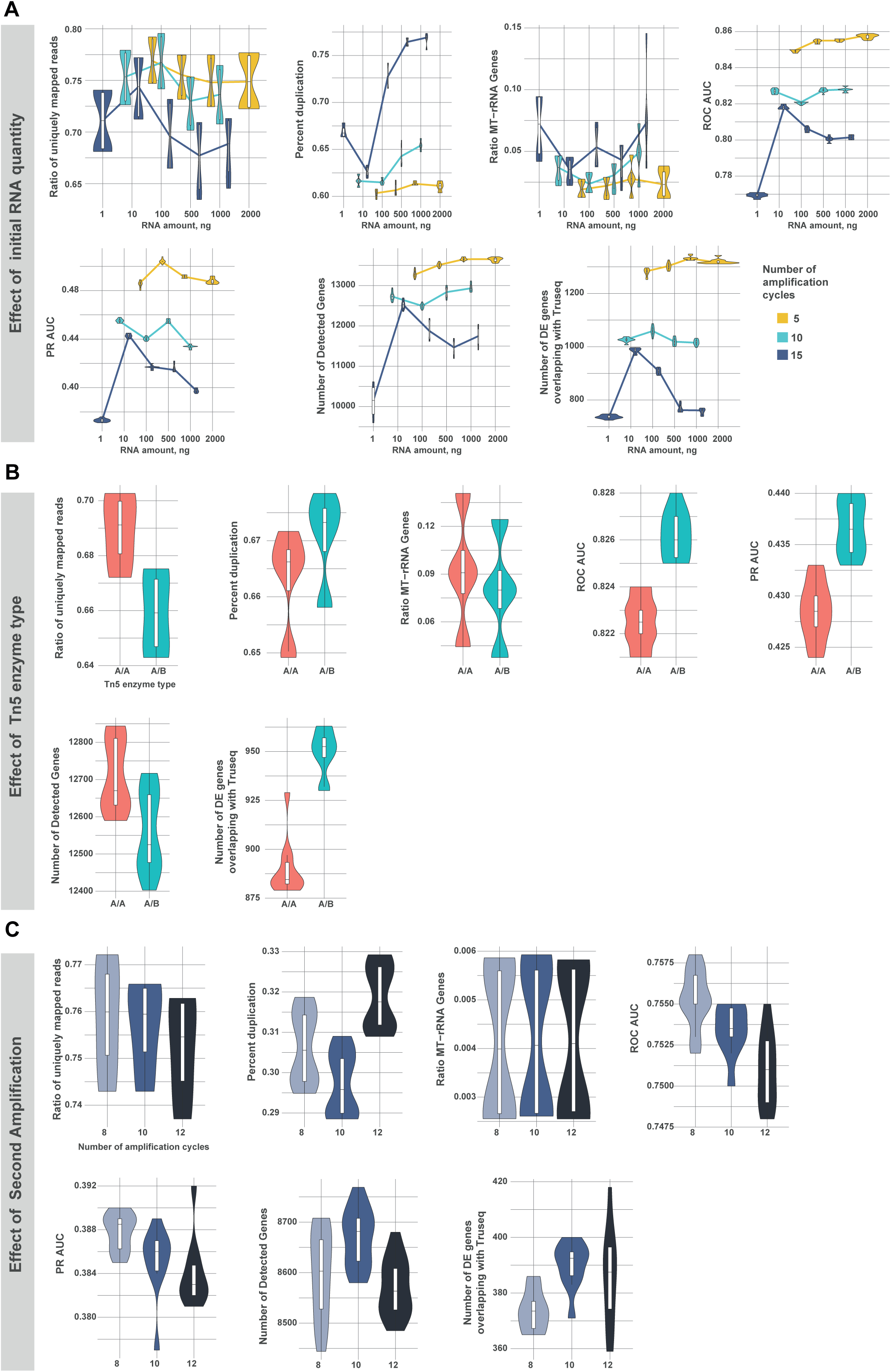
Assessment of the impact of the initial RNA amount (**A.**), Tn5 enzyme type (**B.**) and post-tagmentation amplification (**C.**). Performances were evaluated through variable quality measures: uniquely mapped reads, level of duplication, rate of MT-rRNA reads, ROC AUC (Area Under ROC Curve), precision recall (PR) AUC, number of total detected genes and DE genes overlapping with the “gold standard”. For an unbiased comparison, libraries were downsampled to one million (**A.** and **B.)** or 100k (**C**.) single-end reads (see **Methods**)

**Supp Figure S2.**
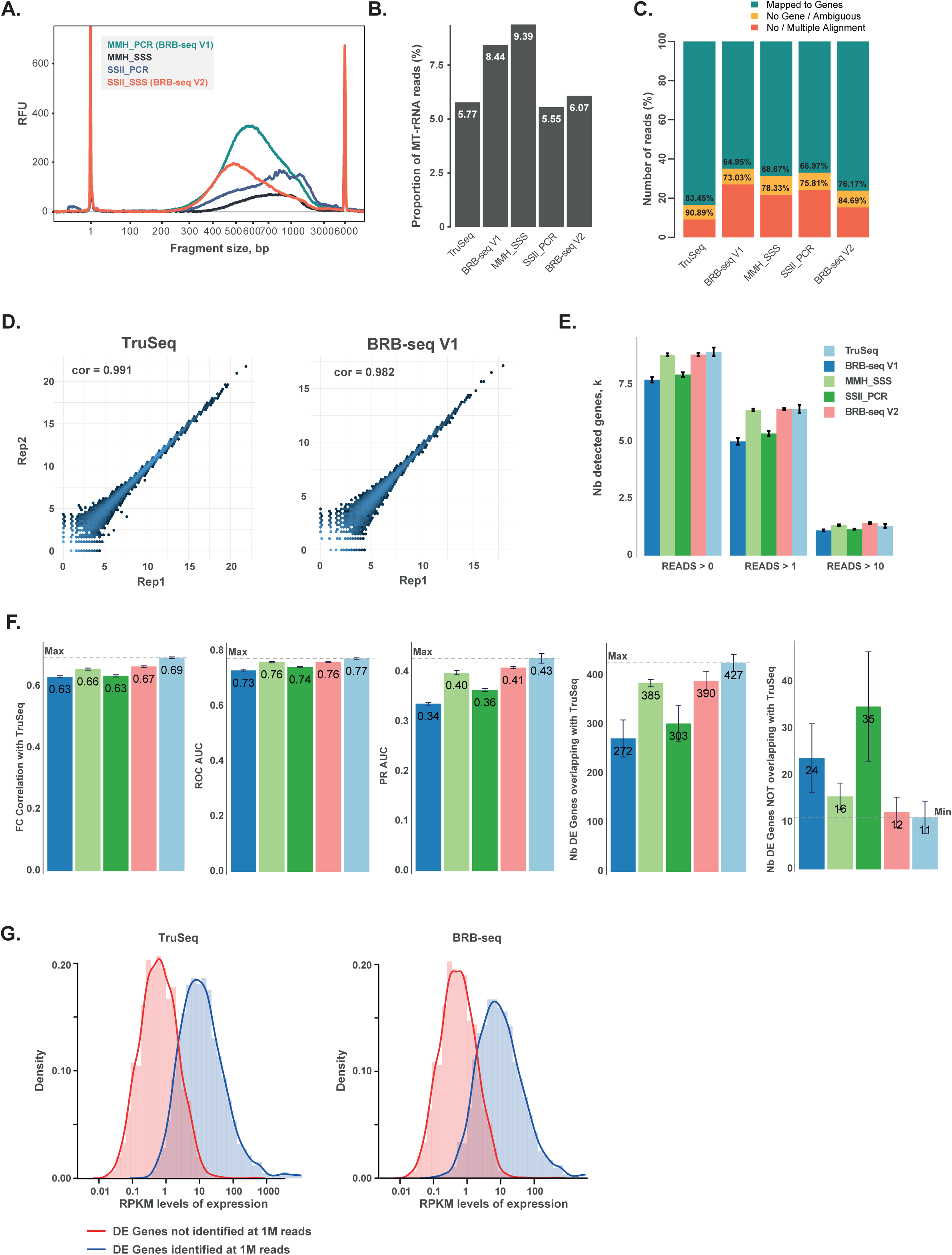
**A.** The profiles of libraries prepared using BRB-seq workflows with variable RT enzymes and second strand generation methods including V1 and V2. The first strand was generated with either Maxima Fermentas Minus H (MMH) or Superscript II (SSII). The second strand was synthetized by either PCR amplification or following the nick translation protocol (SSS). **B.** The percentage of reads assigned to MT-rRNA transcripts per library using variable preparation workflows. **C.** Read alignment performances vs different library preparation workflows. **D.** Correlation of log2 read counts between technical replicates for different protocols (Pearson). **E. & F**. For an unbiased comparison, libraries were downsampled to 100k single-end reads (see **Methods**). **E.** Number of detected genes for TruSeq and different versions of BRB-seq at different cutoffs. For example, ‘Reads >0’ means that a gene is considered detected if it has at least one read for the given sample. **F.** Comparison of TruSeq, and different versions of the BRB-seq workflow, as represented by different quality markers (fold change correlation with TruSeq, Area Under ROC Curve, precision recall (PR) AUC, number of DE genes overlapping and not overlapping with TruSeq). **G.** The distribution of RPKM levels of expression of the DE genes identified (blue) at the sequencing depth of one million reads per sample, that overlaps with the initial TruSeq (paired-end). The curves highlight the distribution of expression level of the genes that are detected (blue) or not detected (red) as DE in the downsampled TruSeq (left panel) or BRB-seq V2 (right panel). The mean expression levels of detected genes (blue, mean RPKM ~10) are higher than those of non-detected ones (red, mean RPKM ~0.5). No visible difference in the detection power was observed between TruSeq and BRB-seq protocols.

**Supp Figure S3.**
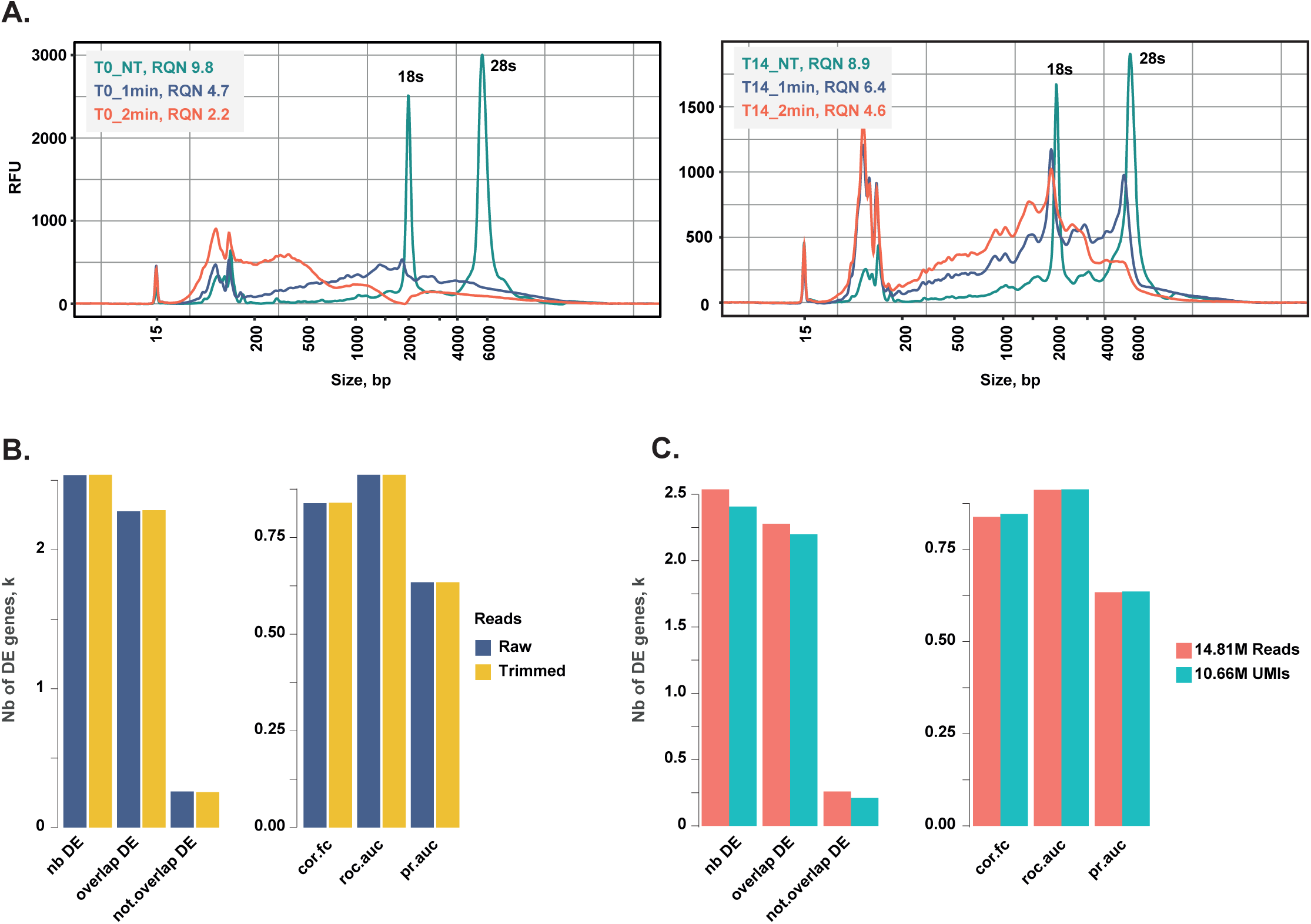
**A.** The RNA profiles of intact samples and their degraded counterparts after one or two minutes of fragmentation. RQN, RNA quality number (maximum is 10). **B.** Impact of read trimming on the data quality for both versions of BRB-seq. **C.** The impact of removal of UMI duplicates on the data quality for both versions of BRB-seq. **B. & C.** The overall performance was assessed according to the number of identified total DE genes (nb DE), overlapping and non overlapping with the “gold standard”; fold change correlation (cor.fc); ROC AUC (roc.auc) and precision recall AUC (pr.auc).

